# Behaviors resulting from the activation of single olfactory receptor neuron class depends on multiple second-order neuron types

**DOI:** 10.1101/2025.05.01.651725

**Authors:** Samuel P. Wechsler, Vikas Bhandawat

## Abstract

Animals rely on olfactory cues to guide critical behaviors such as foraging, mate selection, and predator avoidance. Animals discriminate between different odors because each odor binds to a distinct set of olfactory receptor neurons (ORNs). The relationship between the activated ORN class and resulting behavior is an intensely studied problem. Genetic tools in the *Drosophila* olfactory system make it particularly suitable for understanding this relationship. In this study, we investigate how activity in Or7a-expressing ORNs (Or7a-ORNs) which projects to the DL5 glomerulus, is transformed into aversive behavior. We find that optogenetically activating Or7a-ORNs causes an increase in locomotion speed which results in mild aversion. Surprisingly, silencing the synaptically connected second-order neuron called DL5PN increases the aversion. Silencing DL5PN has no effect on the increase in speed. The increased aversion results from the flies returning to the stimulated area less often. When DL5PN is left intact, flies return more frequently to the stimulated area. Patch-clamp recordings from PNs other than DL5PNs suggest they are activated when Or7a-ORNs are activated. These results suggest that the behavioral effect downstream of a given ORN class is mediated by multiple PN classes. This work advances our understanding of how aversion is encoded and transmitted through early sensory circuits to shape behavior.

## Introduction

Animals, including insects such as *Drosophila melanogaster*, rely on olfactory cues to navigate toward essential resources such as food and mates^10, 76,77^, and to avoid harmful environments, including predators and pathogenic microbes^31^. To meet the challenges of detecting and discriminating between different olfactory cues, animals are endowed with diverse families of odorant receptors ^1,2^ (ORs) that are expressed in olfactory receptor neurons (ORNs) and which bind different odors depending on the ORs they express. In many animals, individual ORNs express only one or few ORs ^3,4^ which determine their response specificity. Thus, ORNs can be classified into a discrete number of classes according to the OR gene^5-8^ they express. ORNs that express a given OR project to a single glomerulus (Fig. 1a) in the antennal lobe (AL), where they connect with the second-order neurons called projection neurons (PNs)^9^. How signals from different ORN classes are processed by downstream circuits to recruit a range of motor programs to facilitate approach, avoidance and other behaviors is intensely studied.

**Figure 1:**
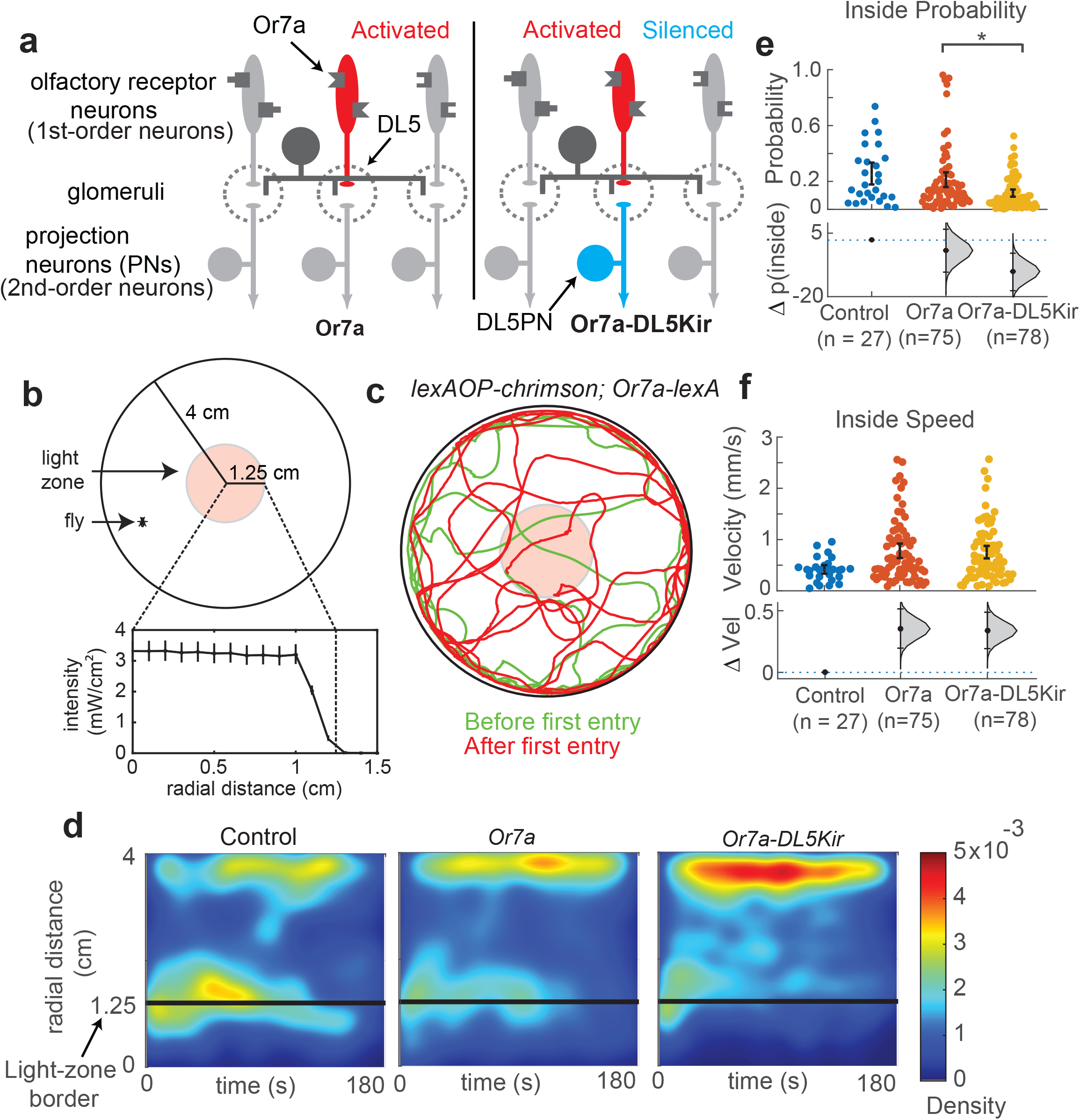
Silencing DL5PN excerbates aversion caused by activation of its pre-syn-aptic ORN, Or7a. **a**. Or7a-ORNs were optogenetically with or without inactivation of DL5PNs. **b**. Schematic of behavioral arena. Light-zone extends 1.25 cm from center. **c**. Example trajectories where Or7a ORNs are activated. **c**. Example tracks before flies first enter light-zone (green) and after first entry (red). **d**. Or7a ORN activation results in a decrease in time spent inside the light-zone which is increased when DL5PNs are silenced. **e**. ORN activation produces a significant increase in speed with or without PN signal silencing, indicating aversion that is not uPN mediated. **f**. Or7a-ORN activation results in an increase in speed which is unaffected by silencing DL5PNs.

A substantial body of work has focused on the motor program underlying attraction^10-12^, both in flight^13,14^ and in walking^15-23^, revealing many conserved mechanisms at play. Similarly, many studies have focused on the relationship between ORN classes activated by a given odor and whether it promoted attraction-associated behaviors ^24-27^. To probe the locomotor mechanism that leads to attraction – i.e., how activities in different ORN classes are related to changes in locomotion – we created a ring arena whose center had a fixed odor concentration, and the periphery did not have any odor^28^. Flies only experienced odors when they went inside the central odor zone; therefore, both the timing and concentration of the odor were known. This study showed that different ORN classes activate many different motor parameters independently^28^, implying that the pattern of ORN activation affected the selection of motor programs underlying the observed locomotion. In subsequent work, we further refined our understanding of the relationship between activated ORN class and behavior using an optogenetic version of the same arena^29^.

In comparison to the study of attraction-associated odors and behaviors, far less is known about aversive behaviors. One reason is that fewer odors and ORN classes mediate aversion^30-36, 48^ compared to those that mediate attraction. Many ORN classes and odors that mediate aversion are considered specialist odors that are processed separately; examples include the processing of CO_2_ and geosmin, a compound that signifies the presence of harmful microbes which flies show an innate avoidance for. As in the case of attraction, motor programs underlying aversion are diverse. One mechanism is an increase in turns orienting away from an aversive stimulus ^34,37^. Another mechanism is increased speed and decreased angular velocity to promote straight, rapid escape away from an aversive stimulus^37^. In the presence of airflow where attractive odors produce an upwind surge behavior, aversive odors can produce freezing^30^. Another mechanism is rapid retreating movements such as backward walking^38^. Additionally, more context-dependent observations can indicate aversion, such as the suppression of oviposition in the presence of an aversive stimulus^30,39^. Much like with attraction, there is limited information on how specific ORN classes influence specific aspects of behavior motor programs.

Compared to the work done to assess the relationship between activity in individual ORN classes and the resulting behavior, much less has been done to assess the relationship between the downstream PNs and behavior. It is widely assumed that behaviors downstream of ORN activation are mediated by the corresponding cognate projection neurons (PNs), the second-order neurons of the fly olfactory system (Fig. 1a). In many studies, the ORN class, the glomerulus it projects to, and the corresponding PN that projects from that glomerulus are used interchangeably. This interchangeability is understandable given that each PN class has been shown to receive inputs from a single, well-segregated neuropil called a glomerulus^40^, and each glomerulus receives inputs from a single ORN class ^41,42^, representing a highly specific information channel connecting ORN ⟶ PN ⟶ higher-order brain centers. However, there are both inhibitory and excitatory interactions between glomeruli^56-61^ making distributed coding at the level of PNs possible even when a single ORN class is activated.

Previous work has shown evidence both for a one-to-one correspondence between ORNs and PNs, and for a more distributed code at the level of PNs. In one such study, aversion was shown to be mediated by a single ORN class^35^. By functionally silencing all ORNs and rescuing a single ORN class, flies showed robust aversion when that rescued ORN class was activated^35^. Surprisingly, this aversion persisted when a large population of PNs including the one downstream of the stimulated ORN was silenced^35^ implying that the downstream PN was not necessary for facilitating this aversive behavior. In contrast, the one-to-one relationship was observed in the case of another ORN class^43^. Activation of Or56a-ORN, which selectively senses the harmful microbe, geosmin, produced a robust, odor-dependent reduction in oviposition^43^. When the cognate PN, DA2-adPN was simultaneously silenced, the suppressed oviposition was abolished.

In this study, we address two issues. First, we relate activity in Or7a-ORNs to motor programs related to aversion. Second, we study whether DL5PNs, the PN downstream of Or7a, are necessary for mediating its role in aversion. Or7a-ORN is a broadly tuned, aversive ORN class that projects to DL5 glomerulus ^32,44-46^. DL5 has a single cognate uPN per hemisphere, DL5-adPN ^44,45^. Or7a-ORNs are traditionally considered aversive, with their activation promoting oviposition suppression ^39,47^ and producing a negative preference index ^31-33,36,47^.

Further, there is only a single DL5-adPN per hemisphere, representing an aversive information channel where the ORN signal is routed through a single cell. DL5-adPNs have also been shown to send direct projections to Av1a1 which is a region of the lateral horn which receives projections from multiple aversive ORN classes ^48^, representing a clear overlap between aversive ORN-mediated signals. In this study, we find that the aversion due to Or7a-ORNs increases when the DL5PN is silenced. One reason for this increase is the more frequent return of the flies when DL5PNs are active. We also show that the activation of Or7a-ORNs results in the indirect activation of other PNs. Overall, this study suggests that even behaviors caused by the activation of a single ORN class recruits multiple PNs.

## Results

### Activating Or7a receptors produces aversion that is exacerbated by inhibiting DL5PN

To characterize the influence of activating an aversive ORN class on locomotion and to determine the role that the cognate uniglomerular PN (DL5PN, See Figure 1a) plays in this transformation, we activated Or7a-ORNs in flies in which the DL5PN functioned normally or in flies in which the DL5PN was silenced (Fig. 1a).

We have previously shown that the activation of a single ORN class can significantly modulate specific aspects of fly behavior, such as speed or the frequency of turning^29^, and because DL5PN receives ∼60% of the total projections from Or7a-ORN^44^, with the next most-targeted recipient receiving only 9%, we expect the removal of the cognate uPN to have a significant impact on behaviors mediated by Or7a-ORNs.

We used the binary expression system *LexA*/*LexAOP*^71^ to express the red light-activated channelrhodopsin, *CsChrimson*, (*Chrimson*)^49^ under the control of the Or7a-ORN promoter to optogenetically activate Or7a-ORNs. Because fly photoreceptors have low sensitivity to long wavelength light, the response to red light itself is minimized. Flies that express *Chrimson* under the control of the Or7a-ORN promoter were placed in the Ring arena ^29^, a small circular arena (8 cm in diameter), whose central region – a circular region 2.5 cm in diameter – had a fixed intensity of light ^50^ (Fig. 1b). As a fly walked into the region with the red light (light-zone), the red light activated Or7a-ORNs and the resulting behavioral change was assessed (Fig. 1c). To selectively silence DL5PN, another binary expression system UAS/Gal4^70^ was used to express *mKir2*.*1*, an inward rectifying potassium channel ^51^, in DL5PN using a split-Gal4 line, *SS32205-Gal4*^*46*^ (Fig. S1b). For brevity, we refer to this experimental manipulation as “Or7a-DL5Kir”.

As in previous studies ^29,50,52^, we first measured the fly’s baseline behavior for a 3 min period during which the central light-zone was off (Fig. 1c, green tracks). We then turned the light on and measured fly behavior for an additional 3 min period for a total of 6 min (Fig. 1c, red tracks). The differences in fly behavior can be assessed by plotting how the fly’s density in different sections of the arena changes as a function of time following their first entry into the light zone (Fig. 1d).

We find that when Or7a-ORNs are activated, flies spend less time inside the light-zone than the control flies which express *Chrimson* in the Or7a-ORNs but are not fed retinal necessary to make Chrimson sensitive to red light; however, the decrease in time spent inside is not significant (Fig. 1e). The decrease in time spent inside is consistent with the significant increase in speed (Fig. 1f), which would indicate an aversive response to the stimulation^53^. These results are consistent with previous work showing that activation of a single ORN class is often not enough to produce a significant change in the distribution of the fly^26-28^ but can change specific locomotor characteristic such as speed^29^.

Unexpectedly, this significant increase in speed while inside the light-zone persists when DL5PN is silenced, with the Or7a-DL5Kir flies showing a significant increase in speed compared to the control that is not significantly distinct from the unperturbed condition (Fig. 1e), which we refer to as “Or7a”. The aversive response not only persists, but is exacerbated following the uPN silencing, with Or7a-DL5Kir flies spending significantly less time inside the light-zone compared to both the Or7a and control conditions (Fig. 1d, e).

### Activation of Or7a-ORNs results in at least two independent motor programs

Other than the time spent inside, there is a change in the time that the fly spends between the light-zone and outer border. This is particularly obvious in the case of Or7a-DL5Kir flies (Fig. 1e). This change might be due to the fact that ORN activity takes some time to decay after the fly exits the light-zone. To tie the response of Or7a-ORNs to the distribution of the flies in the arena, we first characterized how Or7a-ORNs respond to stimulus patterns that flies experience in the Ring arena. To this end, we recorded electrophysiological responses from Or7a-ORNs expressing *Chrimson* (Fig. 2a). The behavioral arena has radial symmetry; the fly’s radial position was used to extract the light intensity it experienced. This measured intensity of light in the arena (Fig. 1b) was then used to generate the stimulus pattern experienced by the fly (Fig. 2b, also Fig. 2-S1). We exposed flies to these stimulus patterns while simultaneously performing single-sensillum recordings^54^. One stimulus pattern is shown in Figure 2b. Single trials showing the response of the fly to this stimulus pattern is shown in Figure 2-S1b. Average PSTH (n = 4) is shown in black in Figure 2b. This provided a description of the responses for Or7a-ORNs during ongoing behavior within the arena. We repeated this process with ten different stimulus patterns to generate a dataset that describes a diverse array of interactions with the light-zone.

**Figure 2:**
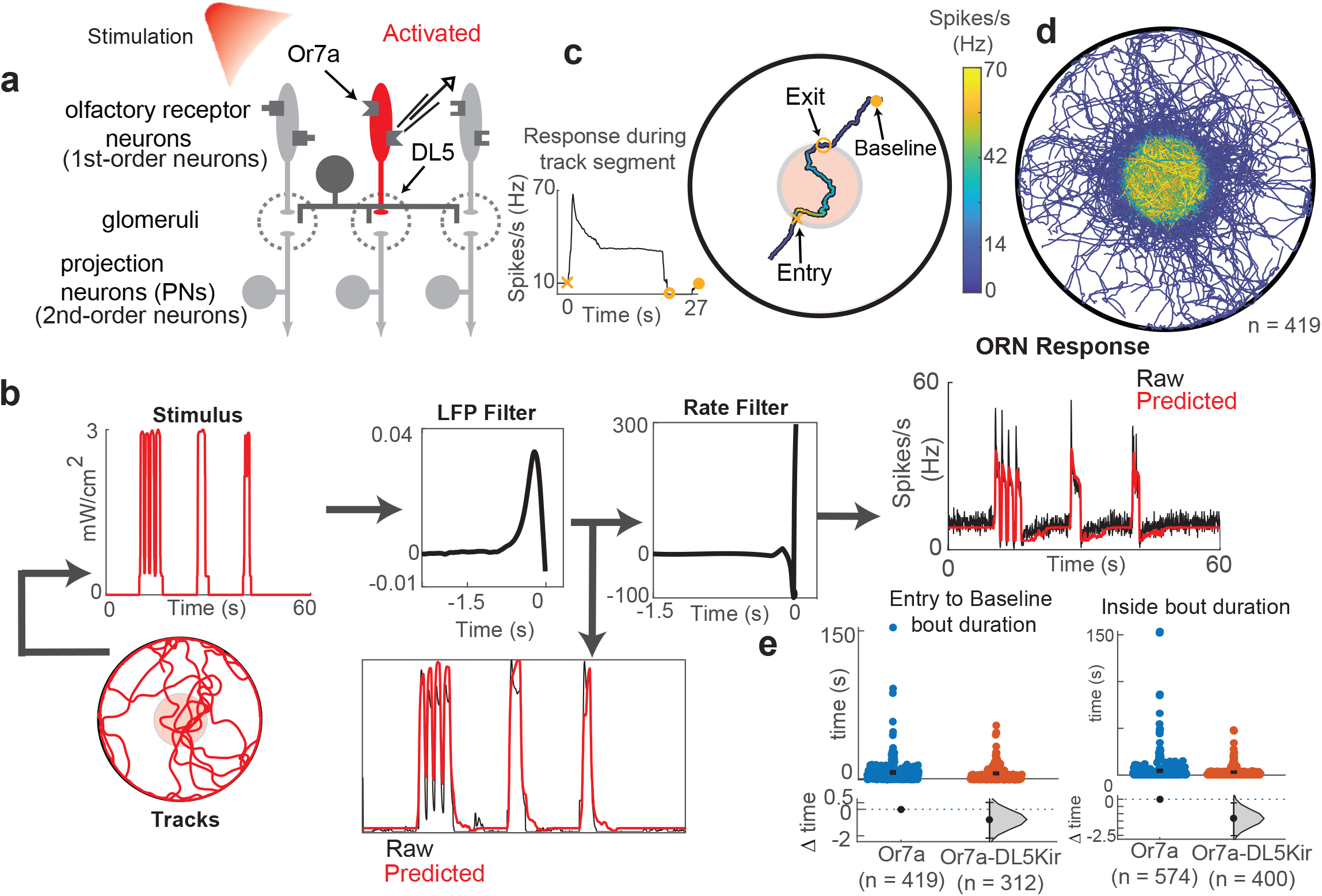
Or7a-DL5Kir flies spend less time in the region where the ORNs are excited and more time in the region in which they are inhibited. **a**. Schematic of experimental setup. Or7a ORN is activated with 617nm light and single sensillum recordings are performed. **b**. Recordings from ten stimulus patterns are used to calculate two linear filters to transform the stimulus to the local field potential and the local field potential to the firing rate. These filters are used to predict ORN response to other stimulus patterns experienced by the fly. **c**. Example entry to baseline track segment. Color represents ORN spike rate. Track segment inside light-zone has a gray outline purely for improved visibility of colors from background. **d**. All entry to baseline events. **e**. Duration of entry to baseline bouts (left) and duration of track segments inside the light-zone (right). There is no significant difference in entry to baseline bout duration when the PN response is silenced. Bout durations while flies are inside the light-zone are significantly shorter when the PN response is silenced.

We used this dataset to construct a neural encoder to predict the responses of Or7a-ORNs to any stimulus pattern reconstructed from fly tracks (Fig. 2b). The neural encoder involved a cascade of two linear filters as previously described^29, 55^ - the first filter described the relationship between stimulus and LFP, and the second filter described the relationship between LFP and firing rate (Fig. 2b). In this way, we reliably modeled the responses of Or7a-ORNs (Fig. 2b). The model’s goodness-of-fit was determined by calculating the coefficient of determination for stimulus patterns that were excluded from the training set; the stimulus pattern in Figure 2b. has an R^2^ value of ∼0.814 (Fig. 2-S1c). This model enabled us to describe the neural response of flies during behavior experiments without additional electrophysiological experiments, and which further enabled us to tie observed changes in locomotion with corresponding response dynamics for Or7a-ORNs.

Using this model, we can infer the ORN activity at any time. Each entry into the light-zone results in a rapid increase in firing rate which adapts before returning to baseline sometime after the fly exits the light-zone. We show an example of a track segment that begins when the fly enters the light-zone and ends when the Or7a-ORNs return to baseline activity (Fig. 2c). The evolution of the neural response during this track segment is shown to the left of the arena diagram, with entry marked with an X, exit marked with an O, and baseline marked with a •. Track segments are color-coded depending on the magnitude of Or7a-ORNs response (Fig. 2c). First, the fly enters the light-zone and experiences a rapid increase in spike rate. This increase is quickly followed by a declining spike rate that is still above baseline while the fly remains inside the light-zone. As the fly exits, Or7a-ORNs do not immediately return to baseline, first experiencing a brief period of inhibition before eventually returning to baseline (Fig. 2c). All track segments for this entry-to-baseline categorization are shown in Figure 2d and indicate that the ORN firing does not return to baseline until sometime after the fly has exited the arena. Immediately following the exit from the light-zone, the Or7a-ORNs are inhibited.

The inhibition of Or7a-ORNs can cause the flies to stay closer to the light-zone. In other words, the flies change their locomotion to stay close to the light-zone. To investigate whether inhibition has a role in influencing fly positioning, we measured the time that a fly spends between entering the light-zone to the time the ORN firing rate reaches baseline (entry-to-baseline). We found that this duration was not significantly different between the Or7a (n=419 bouts across 75 flies) and Or7a-DL5Kir (n=332 bouts across 78 flies) conditions (Fig. 2e, left). The Or7a-DL5Kir flies did spend significantly less time inside the light-zone (Fig. 2e, right). In sum, one consequence of silencing DL5PNs is that flies spend less time inside the light-zone where the ORN firing rate is above baseline, and relatively more time when the Or7a-ORNs are inhibited.

Another difference between Or7a and Or7a-DL5Kir flies is that the Or7a flies return to the light-zone more frequently: 5.5 times for the Or7a flies versus 4.2 times for the Or7a-DL5Kir flies. To characterize the time it takes the Or7a flies to enter the light-zone, we separated the fly tracks into segments that begin when the Or7a-ORNs have returned to baseline and end when the fly enters the light-zone (baseline-to-entry). An example baseline-to-entry bout for Or7a flies is shown in Figure 3a. All extracted track segments for the Or7a flies are shown in Figure 3b. We compare the baseline-to-entry bout durations for the Or7a flies and Or7a-DL5Kir flies (Fig. 3c), which show that Or7a-DL5Kir flies take significantly longer than the Or7a condition to return to the light-zone, with the median bout duration for Or7a flies being ∼8.3s and ∼9.0s for Or7a-DL5Kir flies. When we examined the distribution of baseline-to-entry bout durations, we see that there is, surprisingly, a greater number of bouts < 1 s in the Or7a-DL5Kir flies, but there are greater number of returns for the Or7a flies for most durations thereafter. In particular, the number of returns between 1-4 seconds are particularly high for the Or7a flies. It is possible that the DL5PN response extends beyond the point where Or7a-ORNs have reached baseline, which may explain these differences. To test this hypothesis, we characterized the responses of DL5PNs.

**Figure 3:**
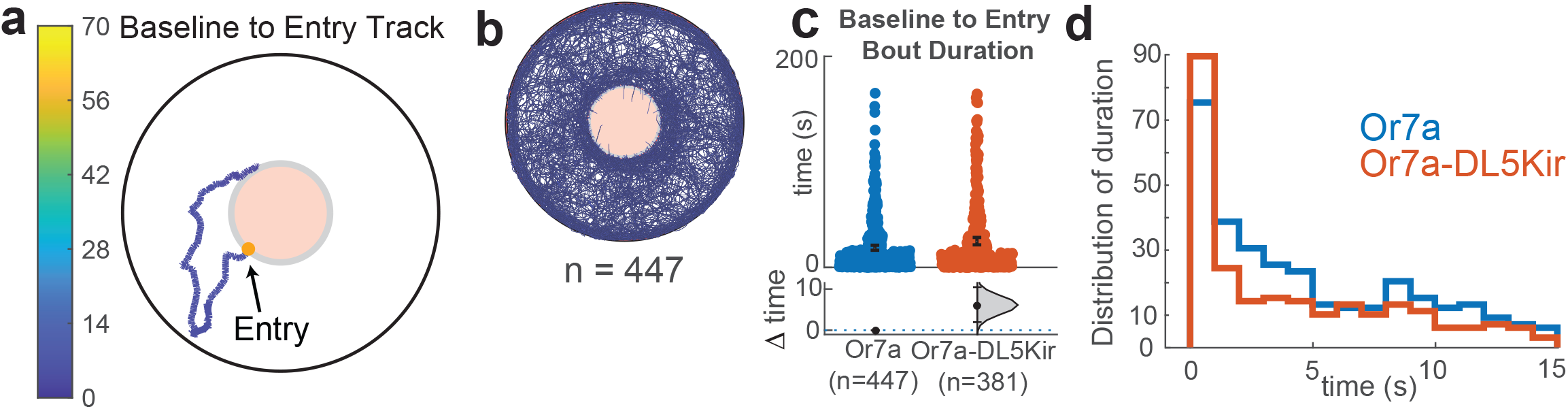
Behavior of the fly during baseline ORN activity is affected by silencing the PN. **a**. Example track of a fly re-entering the light zone after reaching baseline spike activity. Track color is the spike rate. The Entry end point is marked with an orange dot. **b**. All baseline to entry tracks for Or7a flies. **c**. Or7a flies return faster than Or7a-DL5Kir flies. **d**. Distribution of the duration of baseline to entry bouts show that more Or7a-DL5Kir flies returns occur within the first second than the Or7a flies. Thereafter, Or7a flies return more frequently.

### DL5PN activity is important for increased returns to the light-zone

Similar to the method used to characterize Or7a-ORNs response to stimulus patterns experienced during behavior, we measured responses from DL5PN (Fig. 4a) while stimulating Or7a-ORNs with stimuli derived from behavioral tracks. Recording from DL5PN was done as previously described^56, 72^. A rectangular section of the cuticle on the dorsal plane of the fly’s head was removed to expose the antennal lobe. Recordings were directed to DL5PNs by expressing a fluorescent marker. Identity of the recorded neurons was confirmed by filling the cell. The flies were exposed to the same set of stimulus patterns used for the ORN recordings. One stimulus pattern, the corresponding voltage trace for a single trial, raster and PSTH for a single recording is shown in Figure 4b.

**Figure 4:**
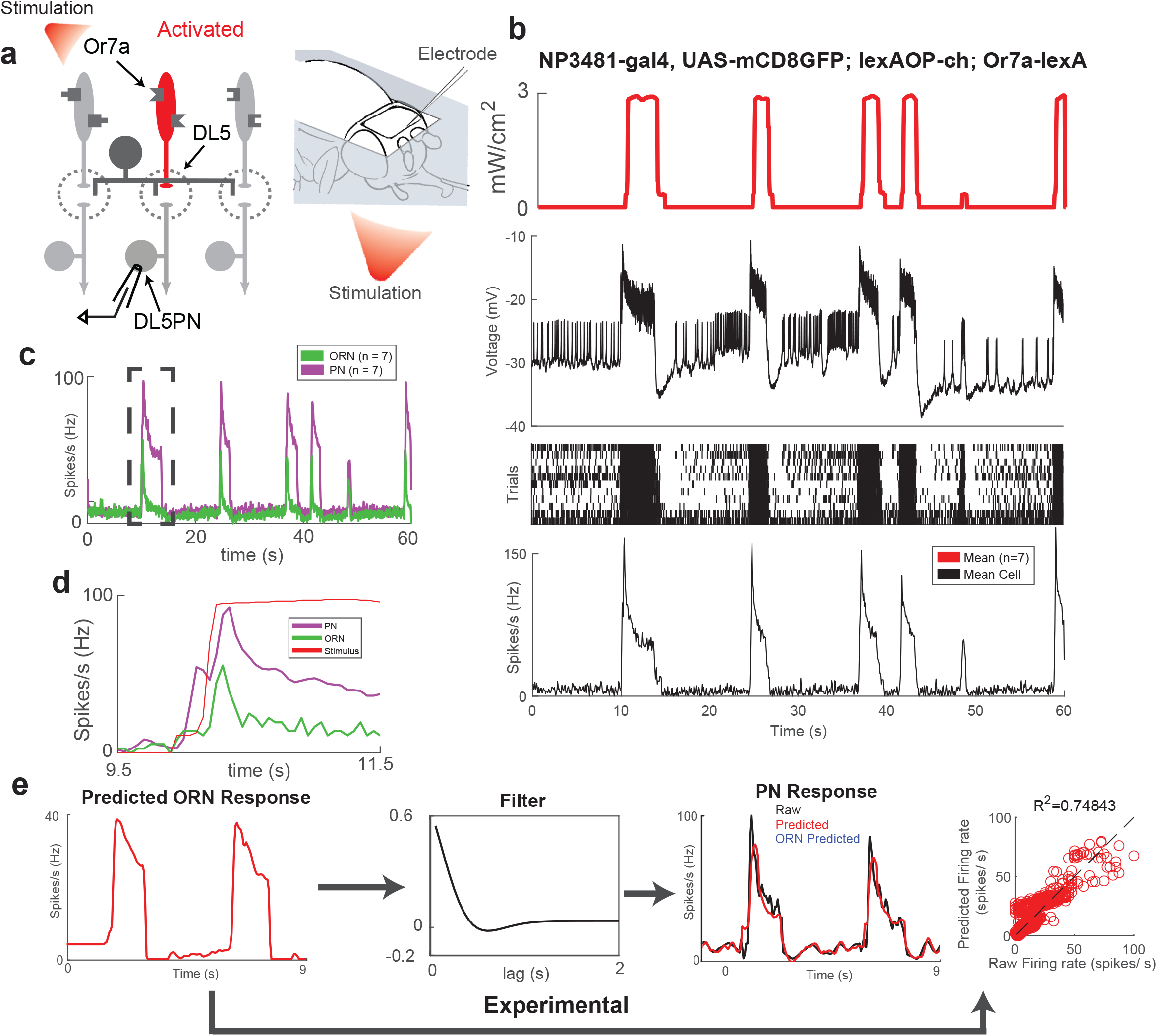
Constructing the response of the PN. **a**. Schematic showing that recording from DL5PN while optogenetically activating Or7a-ORNs. **b**. From the top to bottom: A 60 second light stimulus derived from behavior, sample voltage trace, raster and PSTH. **c**. Mean ORN (green, n=7) and PN (magenta, n=7) response to stimulus pattern in (b). **d**. Close-up of boxed section in (c) showing the dual peak in PN responses. The mean PN response (magenta) shows 2 distinct peaks in spike-rate that coincide with the increasing light intensity when entering the light-zone. **e**. A Gaussian filter was used to transform the ORN firing rate to the PN firing rate.

A comparison between the mean Or7a-ORN response (n=7 sensilla, green) and mean DL5PN response (n=7 cells, magenta) is shown in Figure 4c. DL5PN response is amplified compared to the ORN response, as noted in other studies^72, 75^. We noticed a consistent series of two spikes in the DL5PN response that coincided with the jumps in light intensity as the fly enters the light-zone (Fig. 4d).

Similar to the approach taken with ORNs, we derived a filter to predict DL5PN responses from the predicted ORN response using measured DL5PN response to six stimulus patterns. These stimulus patterns were first used to predict Or7a-ORNs response (Fig. 4e), and this predicted response was then used as the input for a first-order derivative of a Gaussian filter to predict the DL5-adPN response. A comparison between the empirical DL5PN response (black) and predicted DL5-adPN response (red) is shown in Figure 4e (right). The stimulus pattern shown in Figure 4e was not used to train the predictive model. The coefficient of determination for this predicted response was ∼0.75 (Fig. 4e), with the general shape of the response being captured across a series of two stimulus events.

Using the predicted DL5PN response, we first determined whether the DL5PN response persists beyond the time that Or7a-ORNs have returned to baseline. We focused on tracks that are less than 4 seconds, i.e., instances in which flies reenter within 5 seconds. We found that the DL5PN response was at baseline throughout the time the ORNs are at baseline implying that the difference in return time is not due to the activity in DL5PN (Fig. 5a). We next determined the duration of entry-to-baseline and baseline-to-entry bouts based on DL5PN responses. We find that the entry-to-baseline bout durations are not significantly different between the Or7a and Or7a-DL5Kir conditions (Fig. 5b). We also find that the baseline-to-entry durations are significantly longer for the Or7a-DL5Kir condition compared to that of Or7a (Fig. 5c), which is also consistent with the trend observed when using the response of Or7a-ORNs to define the start for baseline-to-entry bouts.

**Figure 5:**
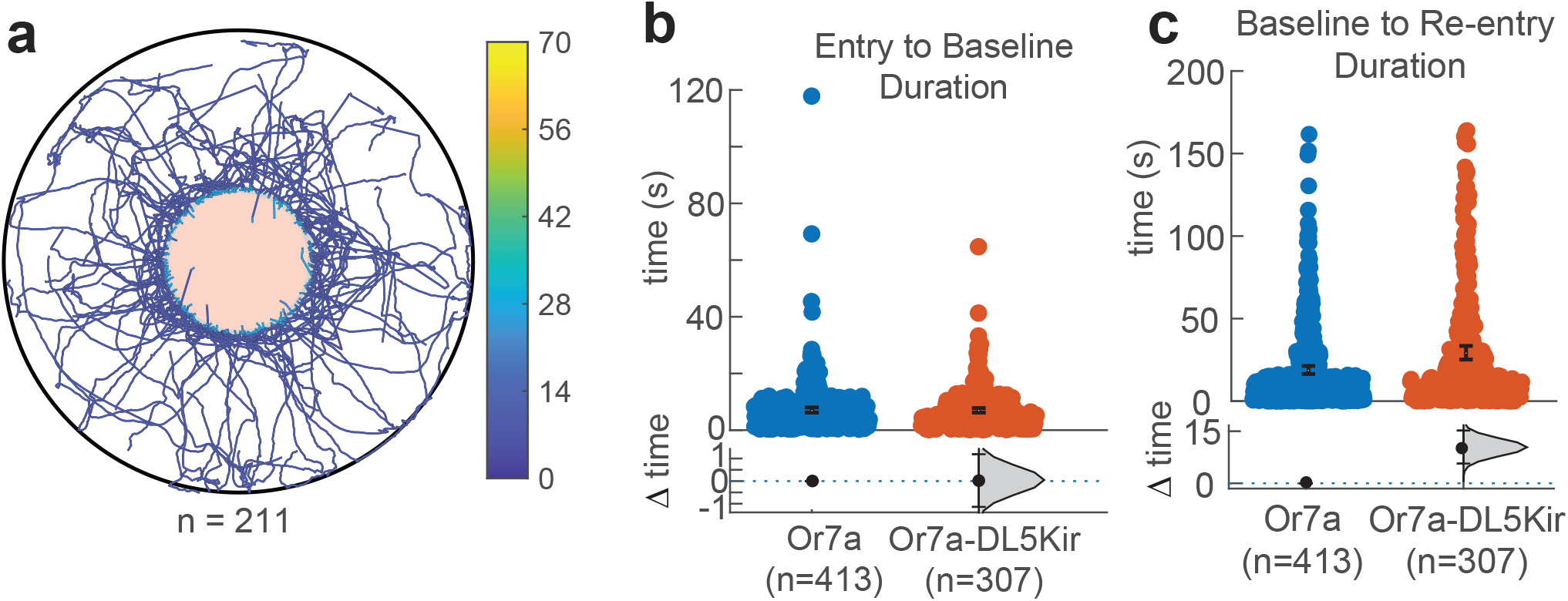
Parsing the fly’s behavior based on PN responses show the same result as dividing it based on ORN response. **a**. Example baseline to re-entry tracks <= 4 seconds in duration. Track onset was based on ORN responses, with track color corresponding to PN firing rate. The PN response does not extend beyond the point where the ORN response reaches baseline. **b**. Entry to baseline bout duration, with track sections ending when the PN response reaches baseline. There is no significant difference between Or7a and Or7a-DL5Kir bout durations. **c**. Baseline to entry bout durations, with track sections beginning when the PN response reaches baseline. Or7a-DL5Kir flies take significantly longer than Or7a flies to return to the light-zone.

Overall, this result implies that one important consequence of silencing the DL5PN is that there is an increase in the time that it takes for the fly to return to the light-zone. However, this increase is not dependent on instantaneous DL5PN activity.

### Activation of Or7a-ORNs drive activity in multiple PNs

It is clear that the effect of Or7a-ORNs on behavior is not mediated entirely by the downstream cognate uPN, DL5PN. Therefore, the activity of Or7a-ORNs must activate other PNs. Or7a-ORNs do not make direct connections to any PNs other than DL5PNs^44^, so any observed activity in these non-cognate uPNs would reflect indirect activation. Indeed, mechanisms for lateral excitation are well-established in the Drosophila antennal lobe^74^. To characterize whether other PNs are activated when Or7a-ORNs are activated, we took advantage of the fact that the lines that we used to record from DL5PNs also label other PNs. We used two lines: NP3481, which labels VM2PN, DM6PN, and VM7PN, and an unidentified mPN (or multi-glomerular PN), and *SS32205* labels one additional glomerulus, DL1. Using these genetic labels, we performed whole-cell patch-clamp recordings on DM6PN, VM7PN, DL1PN, and the mPNs. The identity of each PN was confirmed using biocytin fills.

We find that DL1PNs show moderate responses to the activation of Or7a-ORNs, with an average spike rate of ∼40 Hz (n=3, Fig. 6a). We also find that DM6PNs show weak responses, ∼20 Hz, once the stimulus pattern reached maximum intensity (Fig. 6b). The PN with the most robust response to activation of Or7a-ORNs was VM7d-adPN, with an average spike rate of ∼100 Hz (Fig. 6c), giving it a response magnitude comparable to DL5-adPN (∼100 Hz). We also recorded from three separate, unidentified multi-glomerular PNs (mPNs), which showed weak responsiveness to Or7a-ORNs, ∼25 Hz (Fig. 6d).

**Figure 6:**
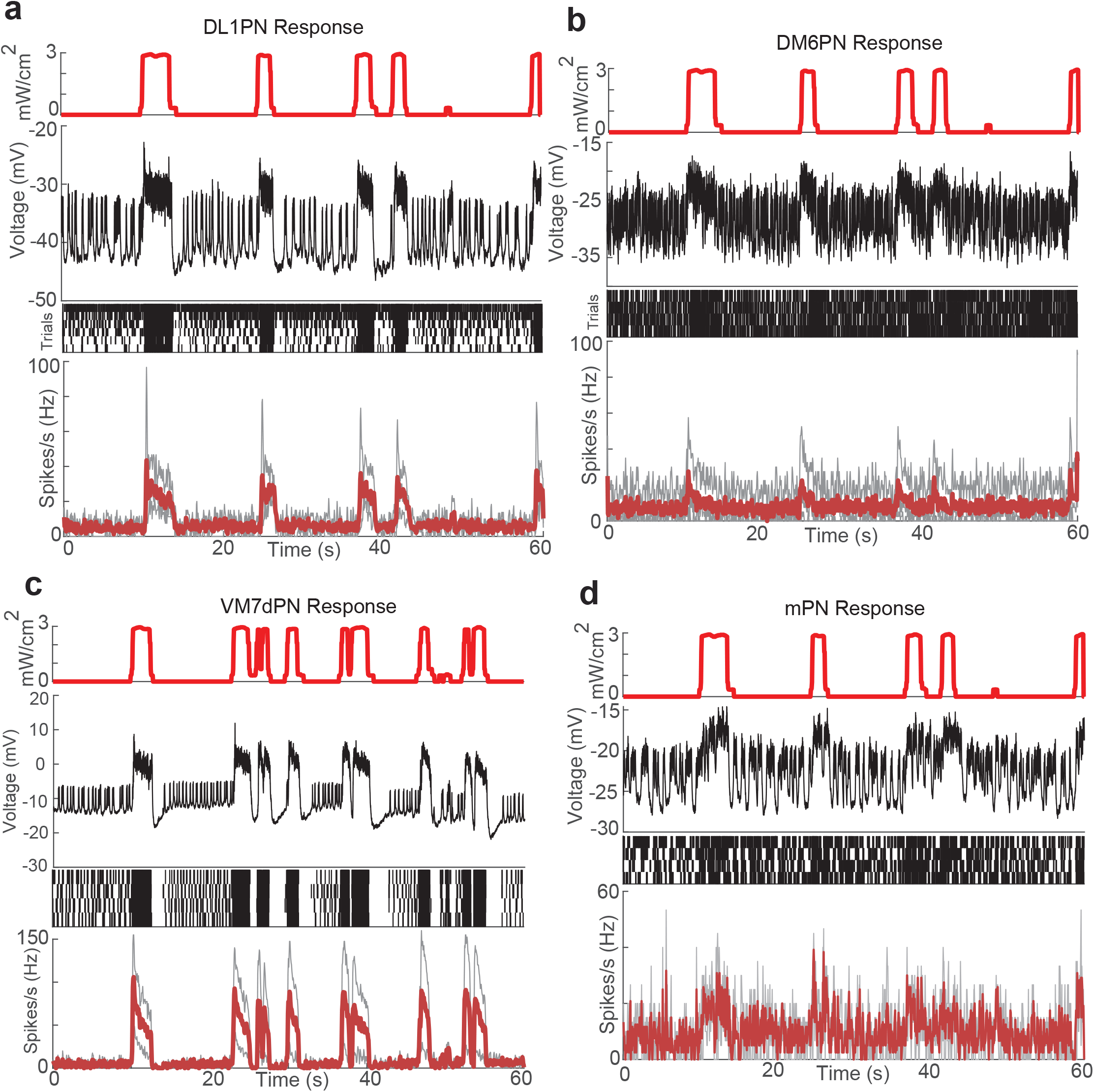
Multiple classes of non-cognate PN respond to activation of Or7a-ORNs. **a**. From the top to bottom: A 60 second light stimulus extracted from fly tracks; voltage trace for a single trial; Raster plot; the average firing rate for each cell (n=3, gray) and mean overall response for DL1PN (red). **b**. Same format as (a). (n=3 cells, gray. Mean DM6PN response in red). **c**. Same format as (a). (n=2 cells, gray. Mean VM7dPN response in red). **d**. Same format as (a). (n=3 separate mPNs, gray. Mean unidentified mPN response in red).

These results show that while DL5-adPNs are the cognate uPN for Or7a-ORNs, the ORN-mediated response spreads through multiple unaffiliated glomeruli via indirect connections and provides a mechanism by which behavioral response persists even when DL5PNs are silenced.

## Discussion

### The control of locomotion by Or7a-ORNs is multi-faceted

In the the case of ORN classes that mediate attraction, a given ORN class mediates changes in different kinematic parameters like speed and curvature^29^, and different ORN class affect different subsets of kinematic parameters, activation of Or7a-ORNs also mediate a change in multiple behaviors. First, activating Or7a-ORNs results in an increase in speed inside the light-zone where the Or7a-ORNs are active. This increase in speed will cause the fly to leave the light-zone faster. The increase observed is strong evidence that Or7a-ORN activation is aversive to the fly as attractive odors or ORNs that mediate attraction typically cause a decrease in speed in the Ring arena. Second, activation of Or7a-ORNs also causes an increased number of returns to the stimulated area. Increased return to the stimulated area counteracts the increase in speed and the net effect is that the time spent inside the stimulated region decreases but this decrease is not significant. This result is consistent with another study that assessed the effect of Or7a-ORNs on locomotion and found that the Or7a-ORNs are neutral^27^. This result is also consistent with previous work showing that activation of a single ORN-class, in most cases, is not sufficient to mediate attraction or aversion^26, 28-29^. Our work also confirms that Or7a-Gal4 does promote aversion and with other ORN class, it is likely to mediate aversion as has been observed in multiple studies which have linked Or7a/DL5 to aversion in studies in which multiple ORNs are activated^24-25^. Overall, this work further supports the idea that the effect of odors on locomotion is not accomplished by a simple binary decision tree: if attractive, go towards. The two effects of Or7a-ORNs on locomotion might reflect the complex role they might play in the fly’s ecology which is still being resolved. Multiple studies have implicated the role of Or7a-ORNs in oviposition, but the effects are different. When Or7a-ORNs are stimulated singly, they have been shown to suppress oviposition^39^. They are also shown to increase oviposition and aggression in combination with food odors.

### Signals from Or7a-ORNs are processed by parallel circuits within the antennal lobe to affect behavior

That the aversion due to the activation of Or7a-ORNs is exacerbated when DL5PN is silenced is surprising. This result implies that the change in speed when Or7a-ORNs are activated is not mediated by DL5PN but mediated by other PNs in the antennal lobe. Consistent with this result, we find that other PNs are also activated when Or7a-ORNs are activated. It is possible that one of the activated PN classes mediates the observed increase in speed. Alternatively, it is possible that the increase in speed results from the combined action of all the PNs, and inactivation of just one PN does not affect the change in speed. It is important to note that another effect of inactivating DL5PN is that the flies spend even less time inside the light-zone. How does the signal spread from the Or7a-ORNs? Based on the connectome^44,57,58^ the only PN that receives direct input from the ORN is DL5PN.The most likely mechanism for the spread of signal across the antennal lobe are connections between ORNs to local neurons (LNs) which can be either excitatory and cholinergic (eLNs) or inhibitory and GABA or glutamatergic (iLNs) ^57,59,60^. A minority of LNs are cholinergic^60^, and when these eLNs are excited, it has been shown that PNs become depolarized^61^. This cholinergic input is not required for excitatory interactions, however, since pharmacological blockage had no effect on the depolarization^62^. These excitatory interactions are mediated through gap junctions^61^.

Past work has predominantly focused on the influence of odor-evoked ORN activation on behavior^78-81^ but odor-evoked inhibition has also been shown to promote aversion^82^, where inhibition of Or85a-ORNs by acetophenone produced a negative preference index, indicating aversion to that odor. The suppression of ORN responses can reduce the spread of lateral interactions, both excitatory and inhibitory, which may promote a distinct motor program in comparison to the motor program recruited by the unsuppressed response. This would suggest that both indirect activation and response suppression may play important roles in mediating ORN class-specific behavior. Our results are consistent with these findings, where silencing the response of DL5PNs modulates behavior differently when compared to that of the unperturbed condition.

In contrast to the observed change in speed, inactivation of DL5PN is sufficient to lengthen the time it takes for the fly to return to the light-zone. Importantly, it is possible that the effect of DL5PN is not mediated solely by DL5PN, as DL5PN activates many multi-glomerular PNs (mPNs). DL5-adPN sends direct connections to 13 mPNs, with a range of 6-50 connections per mPN, and a split of 11 cholinergic mPNs targeted versus 2 GABAergic mPNs targeted ^44^.

Is activation of a single ORN class likely to affect activity in many PNs for other ORN classes as well? The effect of activating an ORN on non-cognate PNs can be both excitatory and inhibitory and it is difficult to predict the overall effect^36^. The effect of activating a single ORN class on the antennal lobe has been measured for two ORN classes. In one study, stimulation with CO_2_, which only affects one ORN class directly, under some conditions resulted in neurotransmitter release from many ORN classes, leading to a distributed representation of CO_2_ ^73^. In contrast, another odor, geosmin, which activates a single ORN class, Or56a-ORN, does not appear to strongly activate any other PN^30^. This question – how signals from a single ORN class is represented at the level of second-order neurons or PNs needs further investigation. Finding odors that activate single ORN classes was a bottleneck. In this study, we use dual binary transcription systems to use optogenetic activation to address this question.

### Hypothesis for the control of behavior at the level of third-order neurons

Silencing DL5PN results in fewer returns to the light-zone. Importantly, this behavior does not depend on instantaneous PN activity and seems to affect the fly’s behavior 1-4 seconds after PN activity has ceased. One hypothesis is that this behavior is driven by the connection of DL5PN to the mushroom body. The DL5PN has strong connections to Kenyon cells in the mushroom body. The cell type most-targeted by DL5-adPN is KCg-m which is involved in short-term memory, and innate responses relating to food or aversive odors ^57,63^ receives 1,548 connections (∼19% of total outputs from DL5-adPN) ^44^.

In regard to DL5’s connection to the lateral horn, DL5PN innervates the same region of the lateral horn as PNs downstream of other ORNs that mediate other aversive behaviors. The lateral horn is an intricately connected neuropil and it is unlikely that the function of a neuron on overall locomotion can be predicted based on its projection pattern alone. It is therefore likely that DL5PN plays a more nuanced role in behavior.

## Methods

### Experimental model and subject details

Flies used in behavior experiments were raised under sparse culture conditions consisting of 50 mL bottles of standard cornmeal media with ∼150 progeny/bottle. Following removal of parents (2 days), active dry yeast was sprinkled in each bottle to enrich larvae’s diet. Bottles were placed in incubators set at 25°C on a 12h dark/light cycle. 10-15 newly eclosed female flies were put in 10mL vials of standard cornmeal media for control experiments, and on food containing all-trans-retinal (0.02% by weight retinal) for optogenetic experiments. All vials were wrapped in aluminum foil to prevent retinal degradation and to keep conditions consistent with the control vials. After 3 days, flies were starved for 24 h inside an empty scintillation vial with half a damp Kimwipe (20 μL of water/ half wipe) for ∼24-32 h prior to experiments. Flies were anesthetized on ice before being placed into the behavioral arena. Fly PN perturbation was performed via expression of human *mKir2*.*1* potassium channel ^51^ in DL5- and DL1-adPNs via *SS32205-Gal4* expression ^46^. Glomerular identity of Kir-expressing PNs was confirmed via co-expression with an RFP tag and visible fluorescent overlap with the ORN (see Figure S1b).

### Behavioral Experiments

Behavioral experiments have been previously described in detail ^29,50^. In brief, experiments were conducted in a 4 cm radius circular arena with a 1.25 cm radius central light-zone. After being placed inside the arena, flies were given 5 min to acclimate before experiment onset. The light-zone was illuminated with red light (617 nm) for the last 3 min of each 6-min experiment. The arena was illuminated with infrared light to enable tracking. Fly locomotion was recorded at 30 frames per second using an infrared video camera (Basler acA20400-90umNIR). Video recordings were compressed to ufmf format prior to tracking^5^. The tracking code models flies as an oval, using the MATLAB regionprops function to extract body orientation and centroid positions. Head position was tracked with the criterion that current head position would be the endpoint along the major axis that makes the smaller turn relative to the previous head position.

### Stimulation within Electrophysiological and Behavioral Arenas

Flies were illuminated using a red (617 nm) light emitting diode (LED) (Thorlabs M617L3) connected to an LED driver (Thorlabs LEDD1B) with the intensity modulated using the driver’s modulation mode which follows the voltage command set with MATLAB. For both electrophysiological and behavioral experiments, LED light was collimated (Thorlabs ACL2520U) and focused with a plano-convex lens (Thorlabs LA1433).

To deliver the same stimulus intensity values in the electrophysiology experiments as measured within the behavioral arena, we first measured the light intensity within the behavioral arena using a photometer (Thorlabs S121C) paired with a 1 mm diameter precision pinhole (Thorlabs P1000D). We then placed the LED at a distance such that the intensity values in the electrophysiological rig mapped to the intensity values in the arena when the driver control voltage values were between 0 and 5 volts. The light was calibrated by applying a series of voltage steps between 0 and 5 volts in 0.5-volt intervals, measuring the intensity with a 1 mm pinhole. We then fit a shifted linear function to map the voltage to intensity and, using these conversions, eleven 60 s behavioral positional trajectories were converted from tracked head position within the behavioral arena to light intensities. These stimulus patterns were up-sampled from 30 Hz to 10 kHz for both single sensillum recordings and whole-cell patch-clamp recordings.

### Single-Sensillum Recording

Single sensillum recording was performed as previously described ^29,64^. *Or7a-lexA > lexAOP-Chrimson* flies were held in a pipette tip using dental wax with the antenna left accessible. The antenna was positioned with glass hooks and visualized using a microscope. A single sensillum was impaled with a saline-filled glass pipette. Responses were passed through a 100x amplifier and filtered with a 5 kHz low pass Bessel filter.

### Whole-Cell Patch-Clamp Recording

Whole-cell patch-clamp recording was performed *in vivo* as previously described ^56^. The saline concentration was based on ^65^ and was as follows (in mM): 140 K-aspartate, 1 KCL, 10 HEPES, 1 EGTA, 0.5 NA_3_GTP (#G8877), and 4 MgATP (#A9187). PN somata were targeted, with one neuron recorded per brain. The morphology of each PN was confirmed post hoc using biocytin-streptavidin, and nc82 histochemistry, as previously described ^56^ with adjustments to the antibody concentrations *(see Immunohistochemistry)*. PN recordings were obtained under visual control using an Olympus BX51WI with infrared optics and a 40X water-immersion objective. Selective targeting of DL5-adPNs was guided by an enhancer trap line that labels specific PNs with GFP, *NP3481-Gal4, UAS-CD8GFP* ^*66*^. The internal pipette solution used (in mM) was as follows ^66^: 140 K-aspartate, 1 KCl, 10 HEPES, 1 EGTA, 0.5 NA_3_GTP (#G8877), 4 MgATP (#A9187) (pH 7.2, osmolarity adjusted to ∼270).

### Immunohistochemistry

To visualize the biocytin-filled neurons and confirm glomerular identity, brains were first fixed with 4% formaldehyde in PBS for 18 min then washed with PBS for 20 min. They were then incubated in a blocking solution of 5% normal goat serum in PBST with 1:20 nc82 antibody and 1:400 chicken α-GFP for 24 hours.

Next, they were washed in PBST for 20 min then incubated in blocking solution with 1:600 Alexa-488 anti-chicken, 1:400 Alexa-633 anti-mouse, and Streptavidin-568 for 24 hours. After incubation, the blocking solution was removed, and the brains were first washed in PBST for 20 min then washed in PBS before being mounted in Vectashield on a 25×75×1mm slide with a 9mm diameter x 0.12 mm depth well and sealed with an 18×18mm glass cover. Fluorescence microscopy was performed on a Zeiss LSM700 using a 63X oil-immersion objective. Biocytin fills and GFP fluorescence were used to confirm the glomerular identity of the PN is consistent with that of the ORN.

#### ANALYSIS METHODS ELECTROPHYSIOLOGY

##### Spike sorting and spike rate analysis

Data collection and spike sorting were performed using a custom MATLAB graphical user interface (GUI). The local field potential (LFP) was found by applying a 300-ms median filter to the signal, then the raw voltage trace was baseline subtracted via subtracting out the LFP. In this study, we recorded from ab4 sensillum, which contains two types of neurons: ab4A-B ^67^. These neurons can be differentiated by their waveform and spike amplitude, and we only considered spikes from ab4A. Spike rate was estimated using kernel smoothing with a 50 ms bandwith ^68^. For PNs, spikes were identified based on peaks in the integral of the baseline-subtracted voltage trace.

##### Filter analysis

A description of generating the filter for predicting ORN spike rate was done as previously described ^29^, with a different set of ten stimulus patterns being used here.

To model the DL5-adPN spike rates, six stimulus patterns were selected, each represented by a single empirical Or7a-ORN and DL5-adPN template response to that stimulus. These responses were low-pass filtered and baseline subtracted. Next, a Gaussian filter and its four derivatives were used as basis functions to fit the ORN-PN relationship. The objective function parameters were optimized using the Nelder–Mead method with MATLAB’s fminsearch function. Using the optimized, first-order derivative of a Gaussian filter and Or7a-ORN responses as the input, DL5-adPN responses were predicted.

#### BEHAVIOR

##### Analysis of the distribution of the fly in the arena

Despite behavioral experiments consisting of 3-min light off, then 3-min light on, flies only start experiencing optogenetic stimulation after entering the central light-zone for the first time. This “first entry” is defined as the first time the fly’s head enters the light-zone (1.25 cm radius circle) after the light turns on. The effect of stimulation on the fly’s distribution within the arena were quantified following first entry.

##### Behaviors were characterized using three methods, as previously described^3^

1. Kernel Density Estimate of spatiotemporal distributions as described above (Fig. 1d). Each fly’s radial head position was aligned based on first entry. Spatiotemporal distributions of this head position were then estimated using MATLAB’s ksdensity function with a Gaussian kernel.
2. Radial Occupancy: The overall probability mass distribution of an average fly being a given radial distance away. Since flies first enter the light zone at different times, the weighted (by relative amount of time after each fly’s first entry) mean and standard error of the mean were calculated. A 2 mm bin size (0.05 radial units) was utilized to generate the distribution.
3. Probability of being inside: This is the proportion of flies inside the central 1.25 cm light-zone as a function of time, smoothed with a 200 ms mean filter.

##### Definition of Entry-to-Baseline and Baseline-to-Entry behavioral states

Fly ORN and PN spike rates do not immediately return to baseline activity after exiting the central light-zone; there is a brief period of inhibition. To account for this continuation of activity beyond the light-zone limits, we define a single interaction with the central light-zone as the point of entry into the light-zone that ends with the first frame (30 Hz) after exiting (radial position > 1.25 cm from center) that returns to baseline activity. Baseline activity for Or7a-ORNs (∼7 Hz) and DL5-adPNs (∼9 Hz) was acquired by taking the avg response for all points where the experienced light intensity was 0. Additionally, we define a single interaction outside the light-zone as the first frame (30 Hz) after reaching baseline that ends with the first frame where the radial distance of a fly’s head reaches <= 1.25 cm from the arena center (inside the light-zone).

##### Estimation Graphics

Instead of calculating p-values, estimation graphics utilize bootstrapping (resampled 10,000 times, bias-corrected, and accelerated) to estimate confidence intervals for either the mean or mean differences. We show individual data points, mean and mean differences using a MATLAB toolbox ^69^.

#### CONTACT FOR REAGENT AND RESOURCE SHARING

Further information and requests for resources and reagents should be directed to and will be fulfilled by the lead contact, Dr. Vikas Bhandawat.

**Table 1:** LIST OF FLY GENOTYPES AND OTHER RESOURCES.

| REAGENT or RESOURCE | SOURCE | IDENTIFIER |
| --- | --- | --- |
| Chemicals, peptides, and recombinant proteins |  |  |
| All trans-Retinal | Sigma Aldrich | R2500 |
| Deposited data |  |  |
| Raw and processed data (behavior and electrophysiology), agent models | This paper |  |
| Experimental models: Organisms/strains |  |  |
| *D. melanogaster*: y[*] w[*] P[w[+mW.hs]=GawB]NP3481 / FM7c | Kyoto Stock Center | DGRC: 113297;<br>FlyBase:<br>FBti0035456 |
| *D. melanogaster*: w[1118]; P[y[+t7.7] w[+mC]=VT033006-p65.AD]attP40/CyO; P[y[+t7.7] w[+mC]=R14A02-GAL4.DBD]attP2/TM6B, Tb[1] | Bloomington Drosophila Stock Center | BDSC: 88840;<br>FlyBase for VT033006:<br>FBti0189894<br>FlyBase for R14A02:<br>FBti0190774 |
| *D. melanogaster*: w[*]; P[y[+t7.7] w[+mC]=UAS-mKir2.1.T2A.tdTomato]attP40/CyO; Pri[1]/TM6B, Tb[1] | Bloomington Drosophila Stock Center | BDSC: 600376;<br>FlyBase:<br>FBti0230568 |
| *D. melanogaster*: NP3481-Gal4, UAS-cD8GFP/ FM7; 13X-ivs-lexAOP2-CsChrimson/Cyo-GFP | Gift from the Dickinson Lab, California Institute of Technology | EJH 306 |
| *D. melanogaster*: NP3481-Gal4, UAS-cD8GFP/ FM6; Or7a-lexA/MKRS | Gift from the Dickinson Lab, California Institute of Technology | EJH 236 |
| Software and algorithms |  |  |
| MATLAB r2020b | MathWorks | RRID: SCR_001622 |
| any2ufmf (part of The Caltech Multiple Walking Fly Tracker) | Branson et al., 2009 | http://ctrax.sourceforge.net/any2ufmf.html |
| Optogenetics arena fly tracker | Tao, Ozarkar, Bhandawat 2020<br>This paper | https://github.com/bhandawatlab/CircularArenaTrackingCode |
| Single sensillum recording and spike sorting GUI | This paper | https://github.com/bhandawatlab/Single-Sensillum-Spike-Sorting-GUI |
| All other software and algorithms | This paper | https://github.com/bhandawatlab/ORN-Optogenetics |

---

**Figure 2-S1.**
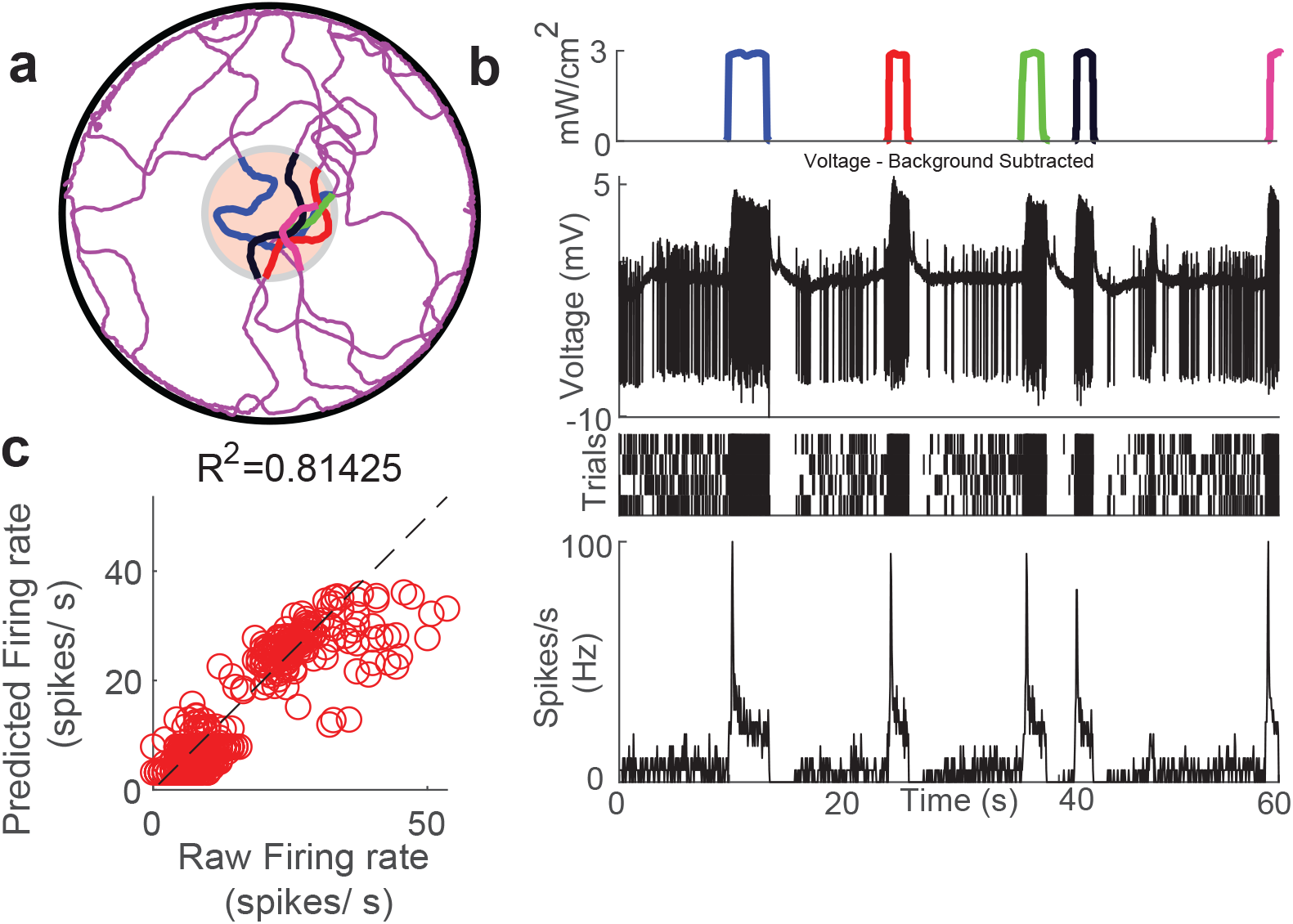
Reconstructing a fly’s sensory experience during behavior. **a**. Reconstruction of fly tracks. **b**. From the top to bottom: A 60 second light stimulus where different colors correspond to different track segments in (a). The distance between stimulus events has been shortened while the light stimulus itself remains unchanged; Local field potential subtracted voltage trace showing spikes; Raster plot of 4 sensilla recordings for the stimulus pattern; the average firing rate of the ORN over the 4 trials. **c**. R^2 calculation between mean Or7a ORN response and predicted ORN response for a stimulus pattern not included in training the model.

## Notes

### Competing Interest Statement

The authors have declared no competing interest.

